# Spatial heterogeneity and microbial terroir: balancing dispersal limitation and cultivar as drivers of microbial diversity in viticulture

**DOI:** 10.1101/2025.08.07.669080

**Authors:** Reid G. Griggs, David A. Mills, Nicholas A. Bokulich

**Affiliations:** Stony Hill Vineyard, St. Helena California, USA; Department of Food Science and Technology, University of California, Davis, California, USA; Laboratory of Food Systems Biotechnology, Institute of Food, Nutrition, and Health, ETH Zurich, Switzerland

**Keywords:** microbial biogeography, microbiome, spatial heterogeneity, wine, marker-gene sequencing, 16S rRNA gene sequencing, internal transcribed spacer, fungal ITS, gamma diversity, kriging

## Abstract

The microbial communities inhabiting grapevines and wines exhibit spatiotemporal patterns linked to region, climate, and cultivar. However, the degree of spatial heterogeneity within and between vineyards and its relationship to cultivar-associated biodiversity selection has not been studied previously. We combined high-density sampling of grapevine microbiota (N = 230) with spatial modeling and satellite imagery in two experiments: (i) two monoclonal Chardonnay vineyards to examine spatial heterogeneity in a genetically homogenous population; and (ii) three old-vine vineyards interplanted with mixed cultivars to investigate the relative effects of spatial distance and cultivar on the microbiota. Contrary to expectations based on monoclonal vineyards, cultivar effects were not apparent in mixed-cultivar vineyards. Instead, we demonstrate extensive spatial variation in the bacterial and fungal communities inhabiting individual grapevines and vineyards, and that community similarity is correlated with spatial distance within and between vineyards. This suggests that dispersal limitation may play an important role in shaping grapevine microbiota, as well as cumulative diversity within the vineyard ecosystem (gamma diversity), with implications for both plant health and wine quality. Spatial models can identify abnormalities in microbial communities, such as contaminant sources within vineyards, and future studies examining microbiota in agricultural settings should account for spatial variation within the study design, e.g., by sufficiently dense spatial sampling or collection of grape musts to avoid undersampling bias. These findings add to the complicated story of microbial biogeography associated with winegrowing and wine quality (microbial *terroir*), highlighting the roles of dispersal and potential microclimate effects in agricultural settings.

## Introduction

Wine and other fermented foods can exhibit profound regional variation in quality characteristics, a phenomenon encompassed by the term *terroir*, which considers regional and site-specific variables (e.g., soil, genotype, climate, and human factors) to be integral components shaping product quality [1]. Substantial evidence suggests that microbiological variation also contributes to this phenomenon in all facets of the winegrowing system, from vineyard soils, to fruit, to musts, and finished wines [2–11]. Regional variation is apparent in both the strain diversity of individual species of yeasts and bacteria [8, 12–15] and in the complex microbial communities that inhabit grapes and musts [2, 3, 9, 10, 16, 17], associated with climate [2, 3, 17], geographic distance [17, 18] and other regional and site-specific factors. Grapevine cultivar is also associated with significant differences in microbiota composition across regions [2, 3, 16, 19–23] and within single vineyards [16], differences that can be linked to specific loci in the grapevine genome, demonstrating that host genotype drives selection of microbial epiphytes [24]. Importantly, variations in the microbial strains and communities inhabiting grapes can be linked to variation in the metabolite composition of wines, indicating the contribution of these microbiota to wine qualities [2, 8, 17]

To date, most studies of vineyard microbiota have examined variation between regions and between vineyards, with only limited investigation of intra-vineyard variation [25]. Previous works have also focused on monovarietal vineyards, i.e., planted to a single cultivar, and such designs complicate the exclusion of block effects from cultivar effects. More information is needed to demonstrate (i) how spatial constraints impact diversity estimates and community assembly within and between vineyards and (ii) how spatial limitation interacts with cultivar effects to impact microbial community composition of grapevines. This could have important implications for microbiota monitoring in agricultural ecosystems, and elucidating the complex interactions between biotic and abiotic factors that shape grapevine microbiota, *terroir*, and wine qualities [26].

To address these questions, two experiments were conducted in parallel to examine (i) the degree of spatial heterogeneity of grapevine microbiota sampled from different clusters within a single grapevine and within the same vineyard, relative to differences between vineyards and (ii) the relative impacts of spatial versus cultivar effects in mixed vineyards. In the first experiment, data were re-analyzed from a previous study [16] of two nearby Chardonnay vineyards in Yolo county, California, that were densely sampled on a single day to examine spatial heterogeneity in monoclonal vineyards; within one of these, a single vine was densely sampled to examine heterogeneity within a single vine (N=12 grapes sampled from different clusters of a single vine). In the second experiment, three old-vine mixed-cultivar vineyards in Sonoma county and geographically close to one another (5.67 km radius) were densely sampled to examine both spatial and cultivar effects in vineyards containing interplanted cultivars. Old-vine vineyards in California are commonly planted to multiple interspersed cultivars, making up Zinfandel-based field blends that are commonly co-fermented. Due to the randomized interplanting of different cultivars, and the common training style, these vineyards present a unique scenario for examining the relationship between microbiota, cultivar, and spatial distance in established vineyards. Results demonstrate a pronounced spatial relationship between and within vineyards, suggesting that spatial dispersion plays a role in microbial colonization of vineyards, potentially explaining the reduced evidence of varietal variation in this work and adding to the developing story of the drivers of microbial terroir in vineyards.

## Materials & Methods

### Experimental Sites

#### Study A: monoclonal vineyard

Data were re-used from a previous study of Chardonnay vineyards at University of California, Davis [16]. These vineyards were selected for dense sampling to examine spatial heterogeneity under monoclonal conditions. The vineyard is cordon trained with vertical shoot positioning, and farmed conventionally. A total of 12 vines each from 5 rows were sampled at regular intervals to conduct a dense sample in a grid. Chardonnay in the Tyree vineyard was planted on Loamy alluvial land and the Reiff soil series, classified as coarse-loamy, mixed non-acid thermic Mollic Xerofluvents. RMI vineyard is planted on the Yolo soil series, described as fine-silty, mixed, superactive, thermic Fluventic Haploxerepts.

#### Study B: mixed-cultivar vineyard

Sampling was conducted in three heritage old-vine vineyards, planted to mixed cultivars. As is common in old California field blends, these cultivars were interplanted, head trained, and spur pruned. The three vineyards were located in close proximity to one another, though were each located in a distinct American Viticultural Area (AVA) within Sonoma County, California, and all vinified at the same winery. Each vineyard contains two dominant cultivars, in addition to much smaller inter-plantings of other cultivars. DEM vineyard is located in Alexander Valley AVA and planted primarily with Zinfandel and Carignan; LEW is located in Dry Creek AVA and planted primarily with Zinfandel and Carignan; and PNZ is located in the Russian River Valley AVA and planted primarily with Zinfandel and Petite Sirah. Data on soil series within blocks was gathered from Soil Web explorer. DEM is planted to the Positas soil series (PsC), a fine montmorillonitic thermic Mollic Palexeralfs in the northern portion of the vineyard, and Manzanita soils series (MbC), a fine-loamy mixed thermic Ultic Palexeralfs in the southern portion of the vineyard. LEW is planted to Arbuckle soil series (AkB), fine-loamy, mixed Typic Haploxeralfs in the eastern portion of the vineyard, and the Pleasanton series (PgB), fine-loamy mixed thermic Mollic Haploxeralfs in the western portion. PNZ is planted mostly to the Haire (HsC) series, a clayey, mixed, thermic Typic Haploxerult, with a small portion of vines planted to the Manzanita series (MbC), a fine loamy mixed, thermic Ultic Palexeralf.

### Sample Collection and DNA extraction

Sampling occurred the night before grapes were harvested, once commercial ripeness was reached. DEM (N=28 samples) was picked on 30 August, 2017; LEW (N=56) on 7 September, 2017; and PNZ (N=50) on 9 September, 2017. Tyree (N=36) and RMI (N=60) were both harvested on a single day (7 August, 2013) when commercial ripeness was reached. Tyree and RMI were both sampled in a regular grid covering the complete block, and avoiding end rows/end vines. Within the old-vine vineyards, fruit samples from representative interplanted cultivars were collected in order to assess the impact of block and cultivar differences at the local scale. Randomized sampling in the vineyard was performed by generating random numbers, and sampling accordingly based on vine number within the vineyard (stratifying by cultivar). Fruit was sampled by aseptically swabbing the entirety of a randomly chosen cluster on each vine with a sterile cotton-tipped swab (Puritan Medical, Guilford, ME, USA). Samples were immediately frozen at -20°C until processing.

DNA was extracted from samples using the Zymo Scientific Fecal/Soil DNA extraction kit (Zymo Research, Irvine, CA, USA) according to the manufacturer’s protocol, with the addition of a 2 min maximum speed bead beater cell lysis step using a FastPrep-24 bead beater (MP Bio, Solon, OH, USA).

### Marker-gene amplicon library preparation and sequencing

Amplification and sequencing were completed as previously described [2]. Specifically, the V4 domain of the 16S rRNA gene was amplified using the universal primer pair F515/R806 [27]. The internal transcribed spacer 1 (ITS1) locus were amplified with the BITS/B58S3 primer pair [28]. Extracted DNA was checked for quality with gel electrophoresis and a NanoDrop spectrophotometer (Thermo Fisher Scientific, DE, USA). Amplicons of the V4 and ITS regions were separately pooled in equimolar ratios, purified using the Qiaquick spin kit (Qiagen), and submitted to the UC Davis Genome Center DNA Technologies Core Facility (Davis, Ca, USA) for library preparation and 250-bp paired-end sequencing on an Illumina MiSeq (Illumina, Inc. CA, USA).

### Bioinformatics Analysis

Sequence data were demultiplexed with QIIME 2 version-2024.10 [29] and denoised with DADA2 [30] to generate ASVs (amplicon sequence variants). Briefly, V4 forward reads were truncated at 155 nts and reverse reads at 165 nt based on quality scores; primers were removed by trimming. Forward and reverse reads were then merged, and chimeras removed using DADA2. ASVs were classified taxonomically using the q2-feature-classifier naive Bayes taxonomic classifier [31] against SILVA SSU (version 13.2 99%) [32] reference sequences prepared and formatted using RESCRIPt [33]. Only forward reads were used in the ITS data to avoid read merging issues, truncated at 220 nt, primers trimmed with cutadapt [34], and ASVs otherwise processed as described for bacterial reads. Fungal taxonomic classification was performed using q2-feature-classifier against the UNITE fungal ITS database (version 10 99%) [35] prepared with RESCRIPt [33]. Any ASVs not assigned at the Class level were removed, as were ASVs accounting for <2 reads, and all negative control samples, and any sequence classified as chloroplast, mitochondria, *Viridiplantae*, *Metazoa*, or 16S rRNA gene sequences classified as *Eukaryota*. This resulted in a V4 dataset with 3,400,287 reads representing 2,881 ASVs in 153 samples and an ITS dataset with 229,641 reads representing 293 ASVs in 105 samples.

### Statistical analysis

Bootstrapped rarefaction and diversity analysis was done using the q2-boots plugin [36] and q2-kmerizer for diversity estimates based on kmer signatures as a pseudo-phylogenetic diversity assessment [37]. Each sample was evenly subsampled at 500 reads per sample, bootstrapping 10 times for resampling with replacement to generate more robust diversity estimates for downstream analysis. Berry microbiota are moderately low biomass and hence the low sampling depth was necessary to retain samples with low read coverage; however, berry microbiota are also relatively low diversity and this sampling depth was confirmed to yield robust and representative diversity estimates via rarefaction.

Alpha diversity metrics (richness and Shannon H) were estimated taking the median of N=10 bootstraps with the q2-boots plugin. Beta diversity metrics (Jaccard distance [38] and Bray-Curtis dissimilarity [39]) were estimated for both ASVs and k-mer tables, taking the medoid of N=10 bootstraps. Principal coordinates analysis (PCoA) was used to visualize these relationships. ANOVA tests (as implemented in q2-longitudinal [40]) and PERMANOVA tests [41] with 999 permutations were used to test for significant differences in alpha and beta diversity, respectively, between groups. Mantel tests with 999 permutations and Spearman correlations were used to test the relationship between spatial distance and bootstrapped beta diversity metrics. Gamma diversity, defined here as the total observed richness of all samples within in a single vineyard, was estimated by measuring cumulative alpha diversity across a given sample set; gamma diversity accumulation curves were calculated by randomly selecting grape samples from within each vineyard (already rarefied at 500 reads per sample) at each given sampling density (*x* samples per site), repeated over 10 iterations to estimate mean diversity and confidence intervals at each density.

Supervised classification was performed using the q2-sample-classifier plugin [42] to train Random Forest classifiers [43] to predict cultivar and vineyard based on microbiota composition. Bacterial and fungal feature tables were rarefied at 500 ASVs per sample, decomposed into k-mers using q2-kmerizer [37], and merged. Random Forest classifiers were trained on these tables using 10-fold nested cross-validation, such that each sample appears in the test set exactly once and accuracy metrics are averaged across all folds. Each classifier consisted of 1000 trees; otherwise default parameters of q2-sample-classifier were used for training and classification.

Spatial interpolation and mapping of microbial diversity estimation was performed using Gaussian Process Regressors as implemented in scikit-learn [44] using a Matérn kernel and 10 optimizer restarts [45]. Spatial mapping was accomplished using cartopy [46], seaborn [47], and matplotlib [48] to plot measured and interpolated values for every individual vine in the vineyard, overlaid on satellite image basemaps derived from Google Tiles via mapping geocoordinates to a plate carrée (equirectangular) projection.

## Results

Extensive within-vineyard sampling was conducted to quantify intra-vineyard variation in microbial community composition and relate this to inter-vineyard variation across local and regional scales. A total of 134 fruit samples were collected from 3 vineyards in Sonoma County, and 96 fruit samples from 2 vineyards in Yolo County (California).

### Microbial community composition heterogeneity in a monocultural vineyard

First we examined the composition of grapevine microbiota in two monovarietal vineyard blocks of *Vitis vinifera* var. Chardonnay in Yolo County to get a “baseline” assessment of how much spatial heterogeneity exists within a single grapevine; within a single vineyard block; and relative to the amount of heterogeneity between two vineyard blocks when cultivar and viticultural practices are uniform.

A single “oversampled vine” (OSV) was sampled 12 times from different clusters to measure within-vine heterogeneity, to compare against the degree of heterogeneity within the same vineyard (Tyree) and between vineyards (i.e., Tyree vs. the nearby RMI vineyard). Results show that within-vineyard beta diversity differences were significantly lower than between-vineyard differences (Figure 1), indicating that both bacterial and fungal communities are more similar between vines within the same vineyard than between vineyards. Moreover, both bacterial and fungal communities were significantly more similar between clusters of the same OSV vine than between Chardonnay vines within the same vineyard. Bacterial ASVs, but not k-mers, demonstrated significantly more similarity between OSV clusters vs. vines within Tyree, suggesting that differences in bacterial communities between vines are driven by subspecies-level differences, but that similar taxonomic groups inhabit the full vineyard. On the other hand, fungal communities demonstrated significant Bray-Curtis dissimilarities in both ASV and k-mer composition, but not Jaccard distance, suggesting that the same sub-species variants could be detected throughout the vineyard, but the abundance of individual variants and different taxonomic groups are more spatially constrained.

**Figure 1.**
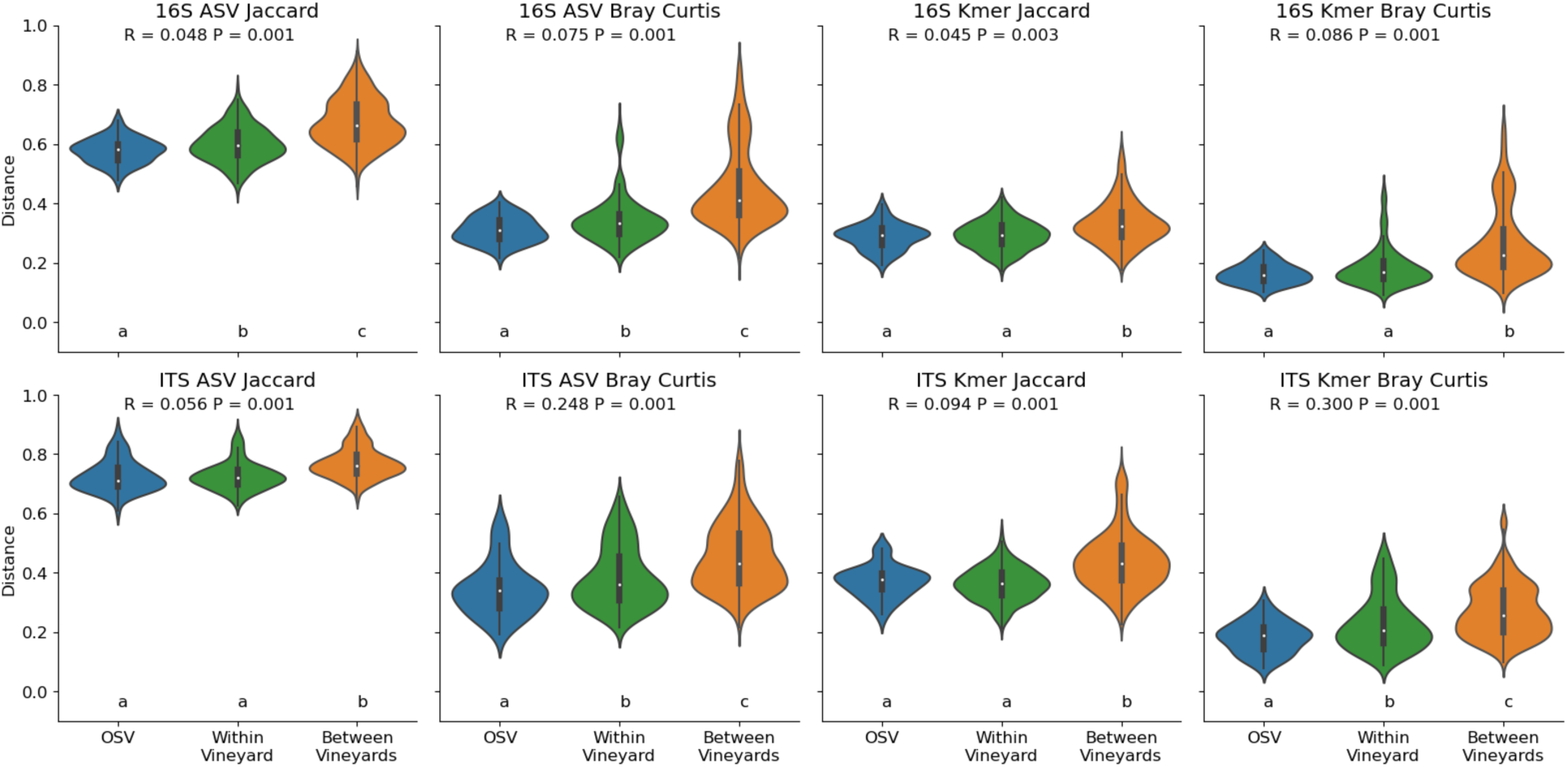
Bacterial 16S rRNA gene (top) and fungal ITS beta diversity (bottom) of Chardonnay grapes varies significantly within a single oversampled vine (OSV, blue), individual grapevines within a single vineyard (green) and between Tyree and RMI vineyards (orange). Bacterial and fungal composition was compared using two beta diversity metrics (Jaccard distance and Bray-Curtis dissimilarity) and two different feature representations (ASVs and k-mer composition, which measures the genetic similarity between each community). PERMANOVA R and P values are indicated for tests comparing the three groups at the top of each panel; pairwise PERMANOVA test results are indicated at the bottom of each panel (groups with different lowercase letters are significantly different).

Significant spatial patterns suggesting dispersal effects were also observed in each vineyard: geodesic distance between individual vines was significantly correlated with beta diversity differences between bacterial and/or fungal communities inhabiting those vines (**Figure 2**). In RMI vineyard, bacterial and fungal alpha diversity (richness and Shannon entropy of ASVs and k-mers) revealed significant spatial patterning (**Figure 3**) correlated with row position in the vineyard (**Figure 4**), further demonstrating that microbial communities vary across a gradient. This effect was not observed in Tyree (P > 0.05), but it should be noted that the vine orientations were roughly North-South in RMI and East-West in Tyree at the time of sampling, suggesting that solar exposition or wind deposition could relate to these differences. Satellite imagery of this vineyard overlaid with spatial interpolation data (**Figure 3**) suggests that local structural features could also explain this spatial effect, with alpha diversity highest near the western edge of the block, which runs alongside an unpaved access road. Together, these results indicate that spatial limitation effects constrain the microbial community composition of grapevines within a single vineyard block, suggesting that either spatial dispersion or localized effects like microclimate gradients, as well as human activities, shape the microbial community structure within vineyards.

**Figure 2.**
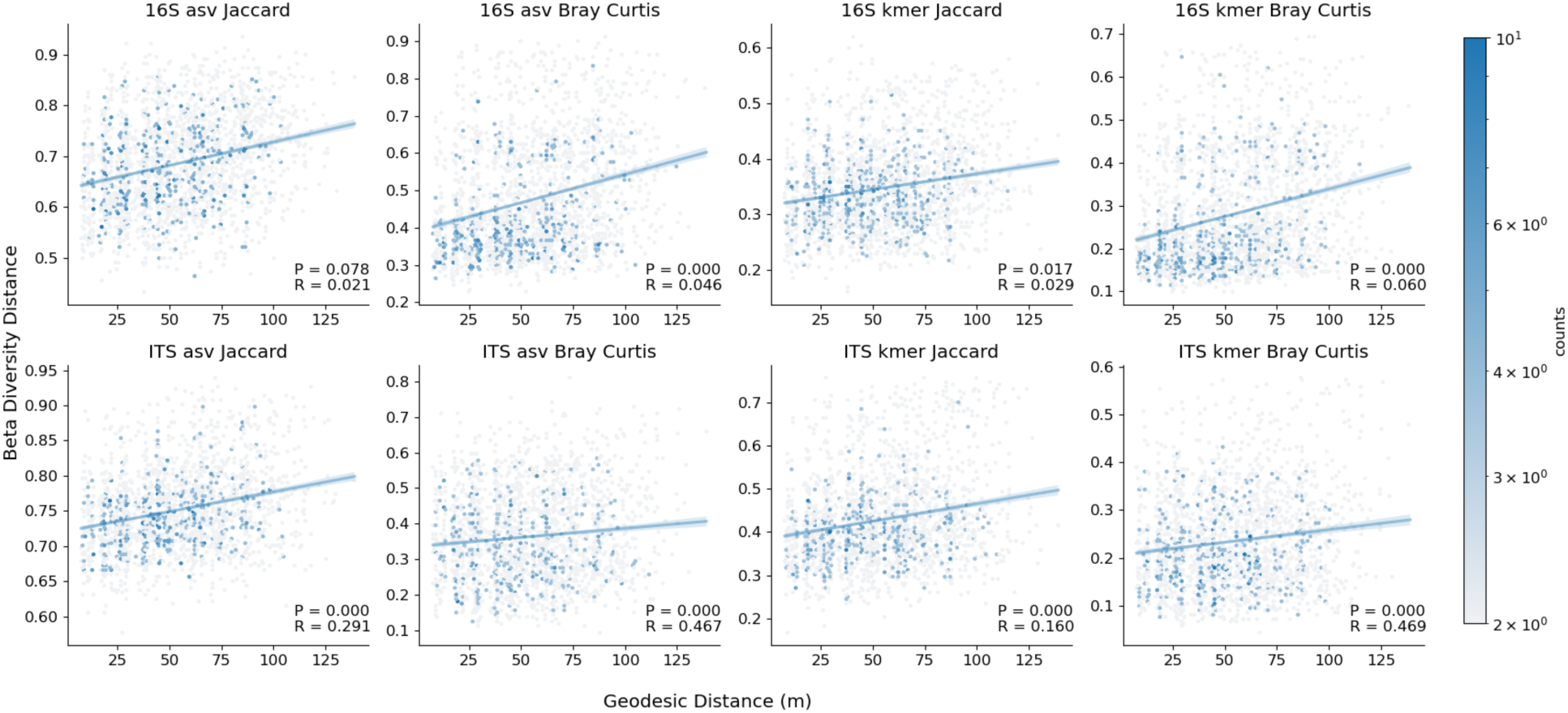
Correlation between spatial distance and beta diversity (distance decay) within single vineyard plots. Bacterial (top) and fungal beta diversity (bottom) pairwise differences between each vine in a single vineyard correlate significantly with geodesic distance between individual vines within the same plot (x-axis), according to different metrics (Jaccard distance, Bray-Curtis dissimilarity) and feature representations (ASVs, k-mers). Pairwise comparisons for both Tyree and RMI are shown. Mantel test results (P-value, Spearman rho) are shown in the bottom-right corner of each panel. These plots are hexbin plots, with the color of each point representing the density of observations at that approximate position. Linear regression lines with 95% confidence intervals (blue shading) are shown for each panel.

**Figure 3.**
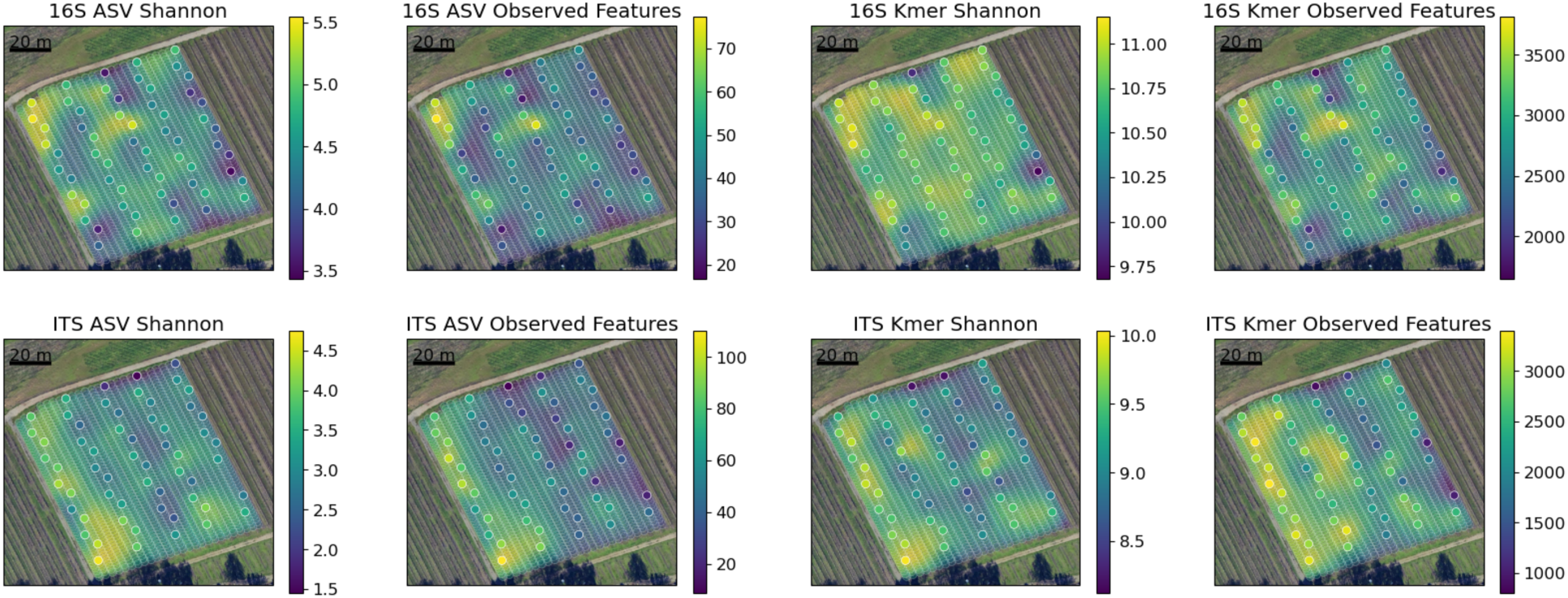
Spatial distribution of bacterial (top) and fungal alpha diversity (bottom) in RMI vineyard. Solid points represent measured values, and semi-translucent points represent predicted values for individual vines in the vineyard using Gaussian process regression for spatial interpolation, overlaid on satellite images of the vineyard.

**Figure 4.**
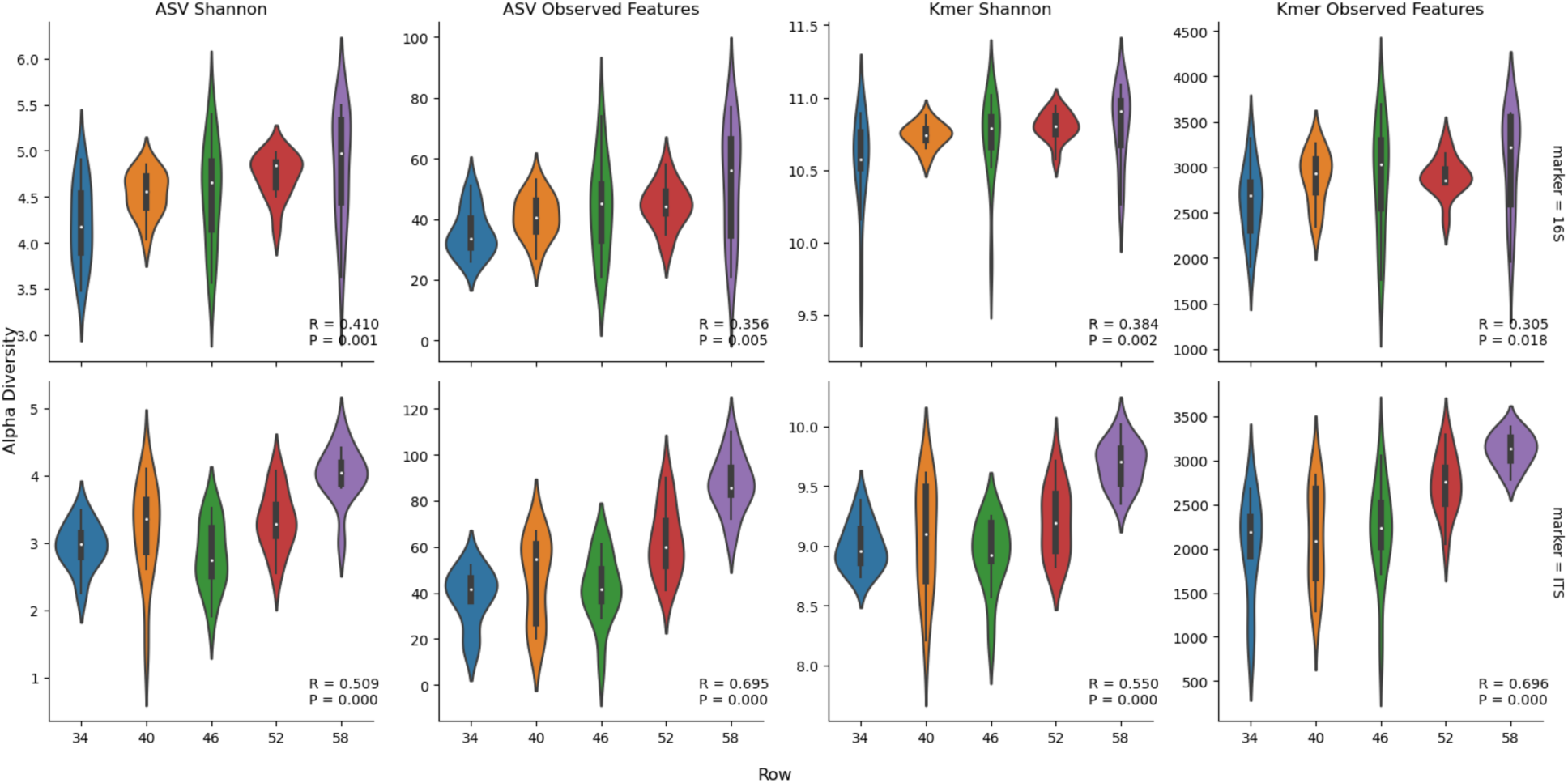
Alpha diversity of bacterial (top) and fungal communities (bottom) vary according to vine row in the RMI vineyard, according to different feature representations (ASVs, k-mers) and metrics (Shannon entropy and observed features, i.e., richness). Spearman rank correlation test results (rho, p-values) indicate the strength of relationship between row position and alpha diversity metrics. Row positions indicate the row number in the entire vineyard, hence row numbers start at 34 (the Easternmost row) and end at 58 (the Westernmost row).

### Spatial effects outweigh cultivar effects in inter-planted vineyards, indicating importance of dispersal limitation for microbiota assembly

Next, we aimed to evaluate the relative impact of spatial versus cultivar effects on vineyard microbiota. Previous studies have shown that grapevine cultivar is associated with grape microbiota composition [3, 16, 23, 49], but these have been in monocultural vineyards making it impossible to parse cultivar from spatial (and in many cases also management) effects. To address this gap, in a separate experiment we examined the microbiota composition of three old-vine vineyards with mixed-cultivar plantings (predominantly Zinfandel), all located within a 5.67 km radius in Sonoma county (California) to see whether site and cultivar effects persisted in this context. As expected, we found that vineyard site was associated with significant differences (P = 0.001) in bacterial and fungal β-diversity according to all feature representations and metrics. Bacterial Bray-Curtis demonstrated the strongest separation (PERMANOVA R_ASV_=0.185, R_kmer_=0.186), indicating that these differences were largely driven by differences in composition, and the presence of unique sub-species variants to a lesser degree (Jaccard distance PERMANOVA R_ASV_=0.042, R_kmer_=0.036); findings mirrored in the fungal communities (Bray-Curtis R_ASV_=0.055, R_kmer_=0.062; Jaccard R_ASV_=0.050, R_kmer_=0.055). Nested Random Forest classifiers also demonstrated highly accurate prediction of vineyard site (micro-average AUC = 0.89, macro-average AUC = 0.90). Alpha diversity also differed between vineyards according to some metrics (Bacterial Shannon entropy P_ANOVA_ = 0.001; Fungal richness P_ANOVA_ = 0.007).

However, cultivar effects were not evident in the mixed-cultivar vineyards in this study. No significant differences (P > 0.05) between cultivars (Zinfandel vs. other cultivars) could be detected in beta diversity by two-way PERMANOVA tests (i.e., after factoring for differences in site) or in alpha diversity by two-way ANOVA tests. Random Forest classification also failed to distinguish between cultivars (macro-average AUC = 0.55). This finding stands in stark contrast to the repeated demonstration of cultivar differences in prior studies of grapevine microbiota, including significant differences in alpha and beta diversity between monocultural blocks under uniform management conditions in the Tyree vineyard [16].

Thus, we hypothesized that spatial dispersion effects could explain the loss of cultivar-specific differences in mixed-cultivar vineyards, due to local transmission of microbiota between neighboring vines. Very high spatial heterogeneity was observed within each individual vineyard (possibly driven by the mixed-cultivar plantings as well as microclimate effects) (**Figure 5**), so no significant spatial effects were observed at the local level (P_Mantel_ > 0.05). However, when comparing the three Sonoma vineyards, significant spatial effects could be observed with some metrics (**Figure 6**). Bacterial Bray Curtis dissimilarity correlated with geospatial distance (Mantel R_ASV_ = 0.177, P_ASV_ < 0.001; R_kmer_ = 0.121, P_kmer_ < 0.001), indicating that the abundance of different bacterial clades varied significantly over moderate geographical distance, but the presence/absence of these groups did not (Jaccard P > 0.05). On the other hand, fungal k-mer diversity was weakly correlated with spatial distance (Mantel R_Jaccard_ = 0.093, P_Jaccard_ = 0.026; R_Bray-Curtis_ = 0.090, P_Bray-Curtis_ = 0.043), but ASV diversity was not by either metric (P > 0.05), suggesting that the presence and abundance of unique genetic variants was spatially constrained, though only to a limited degree.

**Figure 5.**
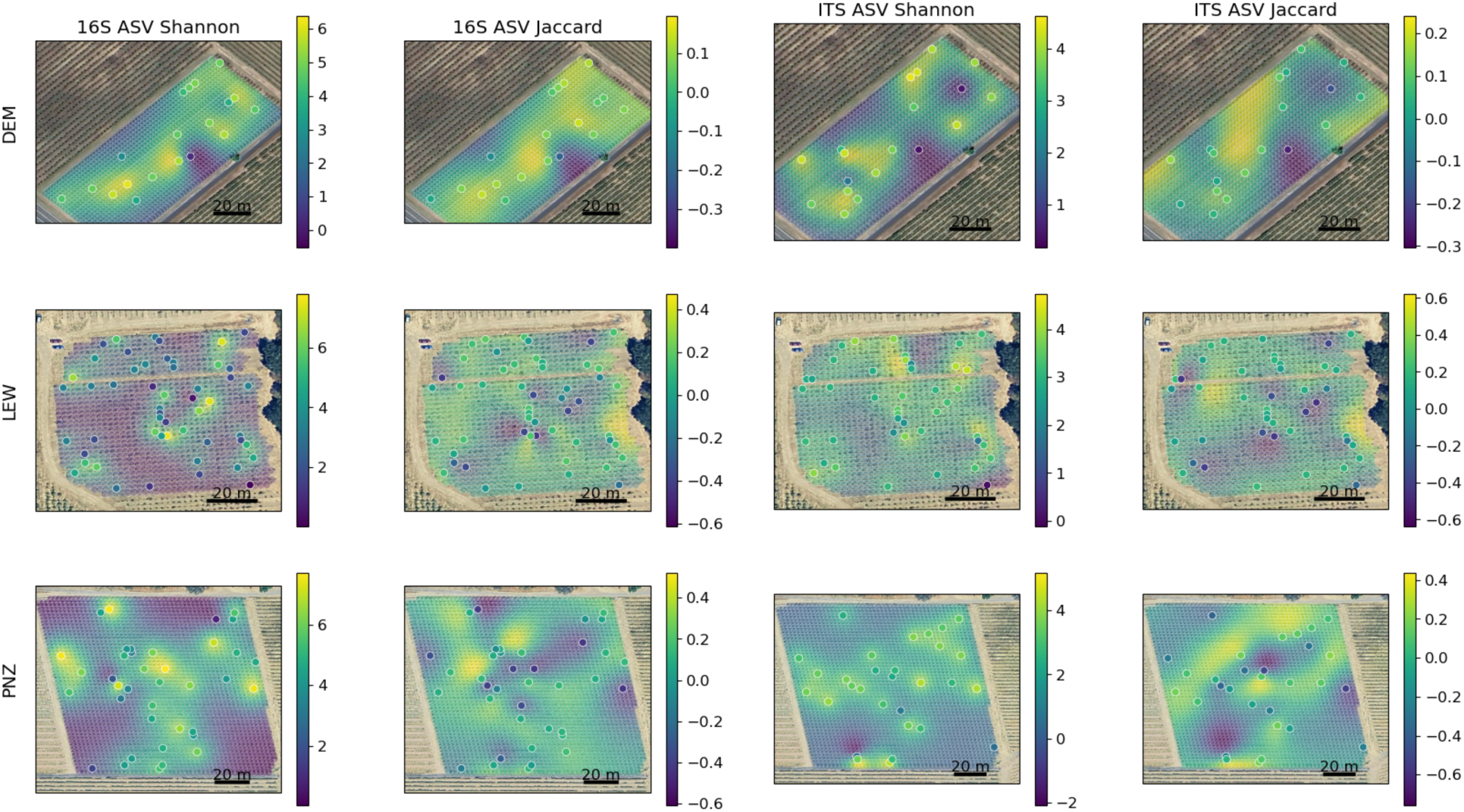
Spatial distribution of bacterial (16S) and fungal (ITS) Shannon entropy (alpha diversity) and Jaccard distance PCoA coordinates in the mixed-cultivar vineyards. Solid points represent measured values, and semi-translucent points represent predicted values for individual vines in the vineyard using Gaussian process regression for spatial interpolation, overlaid on satellite images of the vineyard.

**Figure 6.**
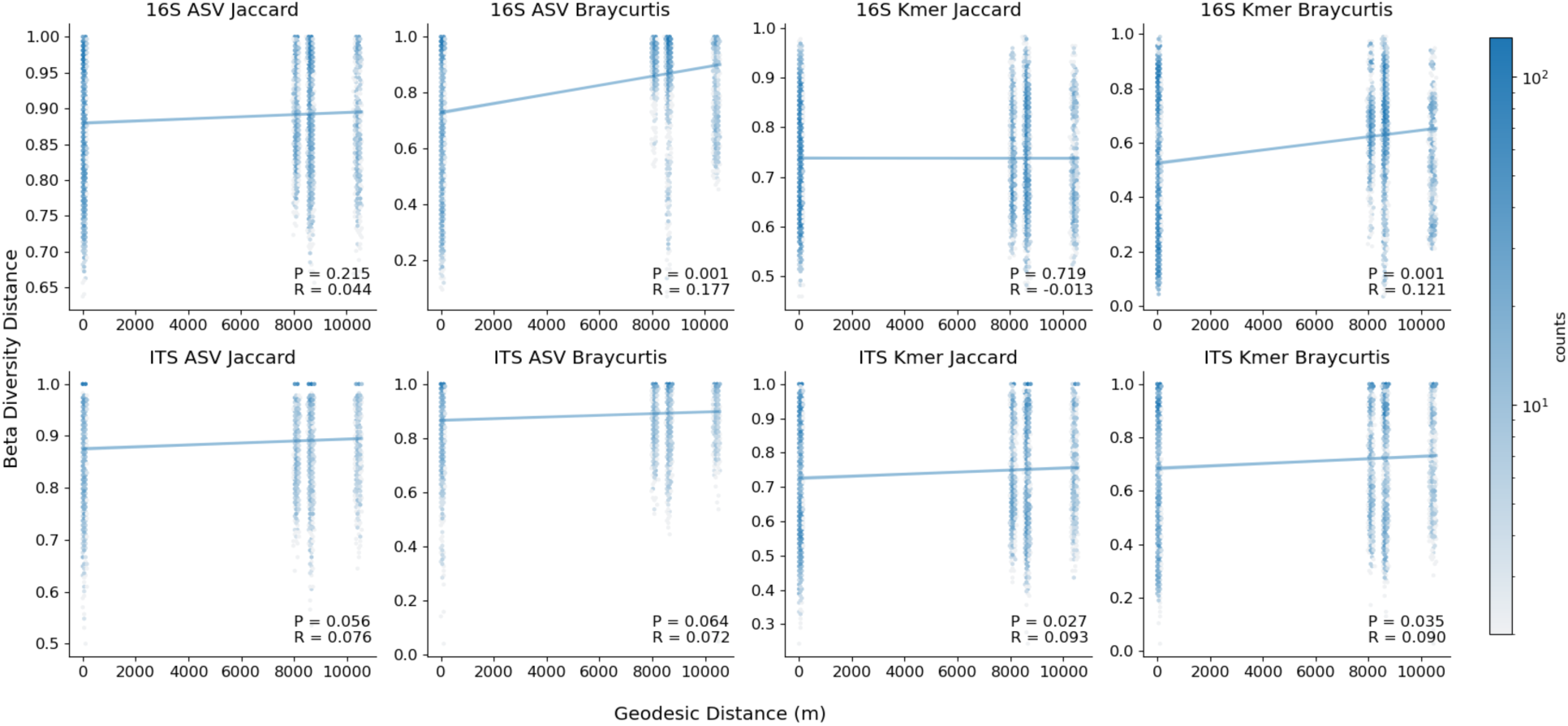
Microbial community composition weakly correlates with geospatial distance between three vineyards in the Sonoma Viticultural Area. Hexbin plots illustrate the relationship between pairwise differences in bacterial (top) and fungal beta diversity (bottom) of each individual vine with geodesic distance (x-axis), according to different metrics (Jaccard distance, Bray-Curtis dissimilarity) and feature representations (ASVs, k-mers). Mantel test results (P-value, Spearman rho) are shown in the bottom-right corner of each panel. These plots are hexbin plots, with the color of each point representing the density of observations at that approximate position. Linear regression lines with 95% confidence intervals (blue shading) are shown for each panel.

### Gamma diversity estimates indicate that spatial heterogeneity contributes to higher biodiversity at the landscape scale, necessitating dense sampling to adequately profile microbial diversity in agricultural plots

The high level of intra-vineyard variation in both alpha and beta diversity observed in this study led us to hypothesize that increased sampling effort would be necessary to adequately estimate the true microbial diversity of a given vineyard, or to detect rare species present at that site. This can be defined as gamma diversity, or the cumulative diversity observed across a landscape unit [50] (in this case, a single vineyard). We performed sample rarefaction, i.e., randomly subsampling samples within a vineyard instead of sequences within a given sample, to determine how sampling effort impacts diversity estimates within each vineyard. As expected, increased sampling density leads to increased measured gamma diversity, eventually approaching the absolute gamma diversity of a site as the sampling effort saturates (**Figure 7**). Bacterial and fungal Shannon entropy fully saturated within ∼10-20 samples in most vineyards, but observed richness did not fully saturate in any of the vineyards examined here. This indicates that detection of the most abundant species could be achieved with 10-20 samples in these vineyard sites, but that an exceedingly large sampling depth and/or sequencing depth would be necessary to detect all species inhabiting a given site. Hence, deep sequencing of must (crushed grape) samples may be a more effective way to estimate total biodiversity in vineyards, as a mixed accumulation of the complete vineyard; this could also explain previous observations of higher mean alpha diversity in must samples than in individual grape samples, in addition to contaminants introduced to the must during grape processing [16]. However, more work is needed to evaluate if very deep sequencing of individual grape samples could also compensate for this effect.

**Figure 7.**
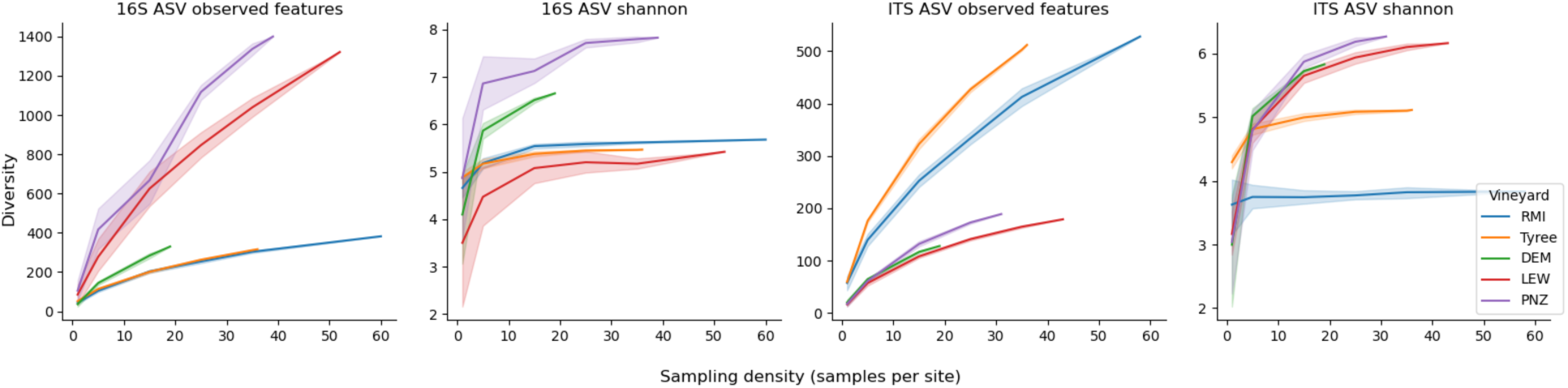
Accumulation curves of bacterial (16S) and fungal (ITS) gamma diversity per vineyard site as a function of sampling density (samples per site, x-axis). At each sampling density, *x* samples per site were randomly selected with replacement over each of 10 iterations, and gamma diversity (Shannon entropy, observed features) estimated across that set. Lines represent mean gamma diversity at density *x*, shading represents 95% confidence intervals.

### Spatial distributions of microbial taxa in vineyards suggest dispersion effects

Spatial mapping of microbiota in vineyards also offers a unique opportunity to examine dispersion of specific taxa within individual sites. As an exploratory approach, Spearman correlations were applied to test for associations between fungal relative abundance, latitude, and longitude of individual grapevines. Several genera — primarily molds — demonstrated spatial relationships, but one stood out as having a strong association in three of the vineyard sites tested here: the phytopathogenic fungal genus, *Erysiphe*, associated with longitude in RMI (R = 0.37) and Tyree (R = 0.67), and with latitude in LEW (R = 0.32). Spatial mapping confirms these trends and shows the spatial relationship (**Figure 8**). *Erysiphe* was detected at the highest relative abundance levels along rows near the edge of the vineyards, highlighting the possibility that dispersion from neighboring fields could be traced using a spatial mapping approach for efforts to trace and control phytopathogens in vineyards. This is intended as a purely exploratory, proof-of-concept demonstration, and further work (including strain typing and absolute quantification) would be necessary to establish true dispersion mechanisms and actual infection risk, goals that are out of scope in the current work.

**Figure 8.**
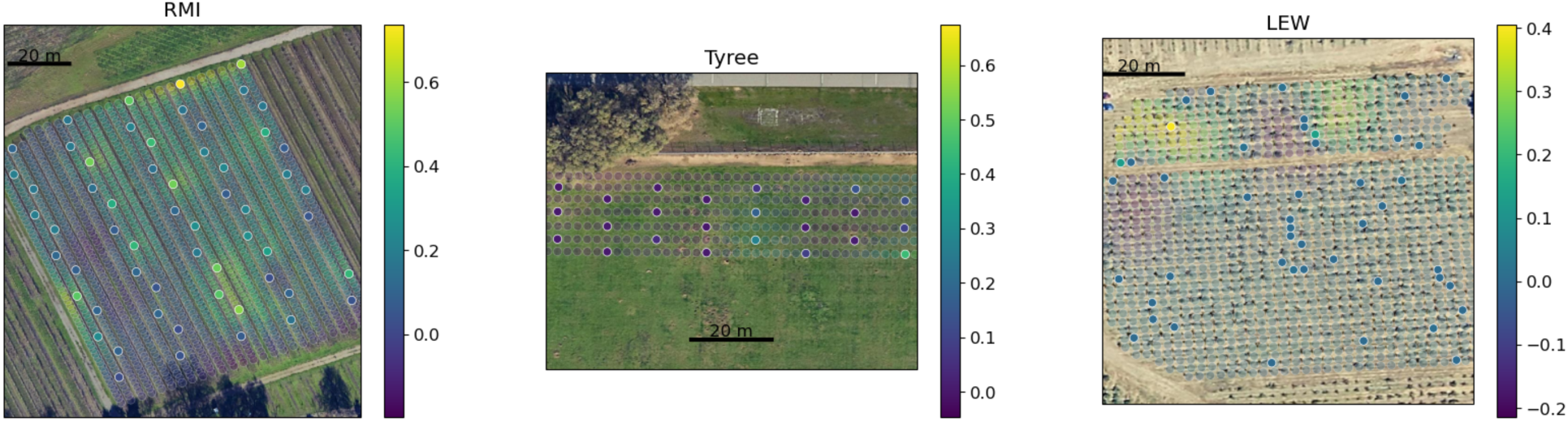
Spatial distribution of the fungal phytopathogen genus *Erysiphe* (powdery mildew) in vineyards. Solid points represent measured values, and semi-translucent points represent predicted values for individual vines in the vineyard using Gaussian process regression for spatial interpolation, overlaid on satellite images of the vineyards. DEM and PNZ are excluded from the plot due to negligible detection levels of *Erysiphe* in those vineyards and insignificant spatial patterns. Scale bars correspond to relative abundance. Note: vines are not visible in the Tyree satellite image, as the satellite image was captured after this vineyard had already been removed.

## Discussion

This work highlights the high degree of spatial heterogeneity inherent in agricultural ecosystems, with implications for plant-microbiome associations in grapevines and other plants. Spatial effects appear to outweigh potential effects of cultivar at the intra-vineyard scale in mixed-vine vineyards, indicating that the plant microbiome is highly sensitive to block effects. In the RMI and Tyree vineyards, a clear distance-decay relationship was observed, suggesting that dispersal limitation shapes microbiota within individual vineyards. This effect was not observed within the old-vine vineyards, suggesting that interplanting of mixed cultivars breaks this pattern due to the secondary influence of cultivar on microbiota selection, though distance-decay relationships were still observed at the regional level, indicating that dispersion effects influence grapevine microbiota at multiple scales.

This study constitutes the first report (to our knowledge) demonstrating such high spatial heterogeneity of grapevine microbiota within individual vineyard blocks, with clear spatial gradients observed in some blocks. Potential explanations for the directional effects of observed diversity (Figures 3-4) include wind or solar exposure, as well as proximity to nearby point sources (e.g., neighboring fields, dirt roads, or other human activities) that shape localized dispersion of microbiota [51]. Alternative explanations include microclimate effects that shape the microbiota directly by influencing growth dynamics, or indirectly by shaping fruit ripening curves, resulting in niche differentiation of the fruit surfaces across the vineyard via changing chemical content of fruit exudates and cell wall structure during ripening [52, 53]. Both mechanisms, differential deposition and niche variability may be at play within vineyards, and future inquiries might attempt to examine intra-block heterogeneity via source-tracking, as well as spatial monitoring of microclimate as well as fruit chemistry.

Whatever the cause, the implications of intra-vineyard microbial diversity in grapevine-associated communities are highly relevant in winemaking, and monitoring microbial diversity in agricultural settings more generally. Grapevine pathogens like *Erysiphe necator* have been demonstrated to be dispersed with regard to prevailing wind direction, training and orientation of the vineyard [54]. Wine-relevant compounds such as phenolics and secondary metabolites like rotundone are also known to vary at the intravineyard scale [55], as do environmental conditions like water holding capacity of soils leading to differential fruit chemistry within a monovarietal block [56]. Thus, winemakers are known to pick fruit with regard to such differences, rather than the arbitrary layout of a vineyard block. Coupling this fact with the presented data here that fruit associated microbial communities might vary at a similar scale, adds a further layer to the collective understanding of the underpinnings of *terroir*. These phenomena likely play a role in creating community-level differences in a similar spatial scale, which could explain spatial heterogeneity in plant health and phenotypic traits, such as metabolite composition, that translate into quality effects in wine and other foods. Modeling intravineyard microbial variation could deepen understanding of the starting materials of wines, and interpolation models might serve as an effective way to test predicted spatial relationships in microbial communities within vineyards.

The cultivar effects observed in this study are unexpectedly subtle, given our previous findings of varietal variation in microbial communities within vineyards (not controlling for block or management effects) [2, 16], across vineyards and regions [3], and known effects of plant genotype on epiphyte communities in grapes [24] and other plants [57, 58]. Balanced with this prior work, it seems likely that genotype effects are nested within regional effects [3, 58–60] and interplanting cultivars in close proximity promotes frequent exchange of microbiota, dispersed by wind, insects, humans, equipment, and other vectors [26], reducing the strength of selection by the plant host that is amplified in monoclonal vineyards. Grapevine genotype clearly impacts the fruit microbiome, shaping epiphyte community assembly via active selection [24]. However, the lack of apparent cultivar effects in mixed-cultivar vineyards suggests that site and block effects are predominant factors shaping the locally available pool of microbiota, and plant genotype acts as a secondary filter. Nevertheless, both effects work synergistically in shaping fruit microbiota within and between vineyards, as mixed-cultivar planting also appears to abolish the clear distance-decay relationship observed in monoclonal vineyards.

Taken together, these findings suggest that cultivar and spatial effects together influence spatial heterogeneity and landscape-scale biodiversity of the plant-associated microbiome in vineyard plots. Host genotype can impact microbiota selection in grapevines [24] and other plants [61], contributing to a legacy effect by which some crops influence microbial biodiversity of the surrounding soil in a phylogeny-dependent fashion [62, 63]. Hence, higher plant diversity is associated with higher microbial diversity in soils [64, 65], other crops [66], and by extension we might expect that mixed plantings of grapevine cultivars as in old-vine vineyards could enhance cumulative microbial biodiversity at the field scale. We do observe that gamma diversity is generally higher in the old-vine vineyards compared to the monoclonal vineyards studied here, but results are confounded by spatial effects; and proper experimental design (e.g., random complete block design of mixed vs. monoclonal plots) will be necessary to properly explore this effect. On the other hand, microclimate and other localized effects (e.g., driven by topographical heterogeneity or management effects) could also lead to higher field-level biodiversity even in monoclonal vineyards.

This leads to the enticing hypothesis that increasing plant biodiversity in vineyards via interplanting as well as use of cover crops may promote higher microbial diversity on grapevines and fruit, leading to higher diversity of microbial communities entering and potentially contributing to wine fermentations. In other words, plant biodiversity in vineyards could promote microbial biodiversity and sensory complexity in winemaking. Interplanting was traditionally practiced in old-vine vineyards to cultivate a resilient system that allows for reliable yield in years where a single cultivar is affected by frost or mildew, or to compensate for quality defects of the dominant cultivar (e.g., interplanting Petite Sirah to increase color, or Carignan for acidity). We speculate that this may also contribute to higher microbial biodiversity as compared with monoclonal vineyards, which could be a double-edged sword depending on the context. On the one hand, interplanting pathogen-susceptible cultivars [67] could introduce a stable reservoir for phytopathogens in that vineyard, potentially increasing risk for the dominant variety and introducing quality defects. On the other hand, interplanting genotypes that select for higher abundance of *Saccharomyces* and other fermentative yeasts on the fruit [24] (e.g., via differences in skin thickness or exudates) could enhance sensory complexity via higher diversity of fermentative organisms in that plot, or even reduce pathogen susceptibility of individual cultivars via increased diversity and colonization resistance in the phyllosphere microbiome. This hypothesis will require extensive validation to explore — via controlled fermentation trials — whether field blends increase microbiological and sensory complexity post-harvest, independent of cultivar chemistry.

The high spatial heterogeneity observed in this study illustrates the challenges of attempting to measure microbial diversity in agricultural settings without key contextual information such as microclimate, soil type, cultivar, and other local variables that can shape heterogeneity in vineyards, with important implications for future studies of plant-associated microbiota in agricultural and natural environments. High spatial heterogeneity is apparent in these vineyards, suggesting dispersion, microclimate, or other highly localized effects. This demonstrates the importance of accounting for spatial heterogeneity in field studies and accounting for spatial covariates when attempting to monitor effects of other treatments on plant microbiota in agricultural settings. Hence, dense sampling is necessary to fully capture biodiversity estimates within such heterogeneous systems, or processing mixed aggregate samples (e.g., grape musts) instead of plant tissues, serving as a sort of “catchment” for examining cumulative biodiversity at the field scale. When studying microbiota *in situ*, fully randomized or grid sampling regimes, as applied here, should be used to evaluate heterogeneity across the field. Gamma rarefaction may be used to evaluate the adequacy of sampling efforts, or to estimate optimal sampling density. Many previous studies have collected only a limited number of samples from different sites, and thus may suffer from undersampling biases that could exaggerate site-specific differences in microbiota profiles (and we would expect, by extension, metabolome profiles as well).

Overall, this study highlights that microbiome assembly in agricultural settings is driven by a complex mixture of biotic and abiotic factors that must be accounted for in biodiversity surveys and assessments of microbial *terroir*. High spatial variation in biodiversity — driven by microclimate, plant biodiversity, and other effects at the vineyard scale — could also promote higher microbial biodiversity in wine fermentations, potentially influencing sensory complexity (as well as spoilage risks) in winemaking. Future studies are needed to evaluate the specific conditions driving spatial heterogeneity in microbial communities at the intra-block level as well as between vineyards and to validate the hypothesis posed here that higher gamma diversity in vineyards could lead to higher microbial biodiversity and sensory complexity in wine fermentations. These findings shed further light on the complicated blend of biotic and abiotic factors that shape microbial biodiversity within and between vineyards, as a key feature of grapevine health and wine quality.

## Data Availability

Data from Tyree and RMI vineyards are deposited in ENA under study accessions PRJEB88146 (ITS sequence data) and PRJEB88181 (16S rRNA gene sequence data).

## Acknowledgments

The authors acknowledge Alice Yu for assistance with sample processing, Zach Stocksdale for assistance with sample collection, and the winegrowers who generously allowed access to their vineyards and supplied ampelographic maps to support this study.

## Author contributions

RGG, DAM, and NAB conceived and designed the study. RGG collected samples. NAB and RGG analyzed the data. NAB and RGG wrote the manuscript. All authors reviewed and edited the manuscript. DAM and NAB acquired funding for this work.

